# B-SMART-Former: An Explainable Transformer-Based Deep Learning Model for Predicting Drug–Drug Interactions Between Biotech and Small-Molecule Drugs

**DOI:** 10.64898/2026.07.23.740240

**Authors:** Fatemeh Nasiri, Mohsen Hooshmand, Mahdi Nouroozi

## Abstract

Drug—drug interactions between biotech and small-molecule drugs play a critical role in medication safety and therapeutic efficacy. However, most existing computational DDI prediction methods focus primarily on interactions between small-molecule drugs, leaving biotech–small-molecule interactions comparatively underexplored. In this study, we propose B-SMART-Former, an explainable deep learning framework for predicting interaction types between biotech and small-molecule drugs. The proposed framework integrates ChemBERTa embeddings and Morgan molecular fingerprints for small molecules with ProtBERT embeddings for biotech drugs, eliminating the need for similarity-based features while leveraging complementary molecular representations. These multimodal features are processed by a hybrid architecture that combines Transformer-based self-attention, residual convolutional learning, and a multi-layer perceptron classifier to capture both global contextual dependencies and local discriminative patterns. The model is formulated as a multi-class classification task and evaluated using stratified 10-fold cross-validation. To improve model transparency, Integrated Gradients is employed as a post-hoc explainability method to identify the molecular features that contribute most strongly to each prediction. Experimental results demonstrate that B-SMART-Former achieves a micro-averaged AUROC of 0.9978 and an AUPR of 0.9682 while relying solely on intrinsic molecular representations, remaining competitive with similarity-based approaches. The proposed framework offers an effective and explainable solution for biotech–small-molecule DDI prediction and provides a practical foundation for future computational drug interaction studies.

## 2 Introduction

The rapid advancement of modern pharmacotherapy has substantially improved the prevention and treatment of numerous diseases, leading to increased life expectancy and better clinical outcomes [1]. However, the widespread use of multiple medications has also introduced significant challenges in ensuring medication safety. Polypharmacy has become increasingly prevalent, particularly among elderly individuals and patients with chronic conditions such as cancer, cardiovascular diseases, and diabetes, who often require the concurrent administration of multiple therapeutic agents [2].

The simultaneous use of multiple drugs substantially increases the risk of drug–drug interactions (DDIs), in which one drug alters the pharmacological effect of another. Clinically significant DDIs may reduce therapeutic efficacy, increase drug toxicity, trigger adverse drug reactions (ADRs), prolong hospitalization, and, in severe cases, result in life-threatening complications or death [3–5]. Consequently, the early identification of potential DDIs has become an essential component of drug development, regulatory assessment, and clinical decision-making.

In recent years, biologic therapeutics, including monoclonal antibodies, recombinant proteins, peptide-based drugs, and vaccines, have become indispensable for treating complex diseases such as cancer, autoimmune disorders, and inflammatory conditions [6, 7]. In clinical practice, these biologic agents are frequently co-administered with conventional small-molecule drugs to enhance therapeutic outcomes. While such combination therapies offer considerable clinical benefits, they also introduce complex interaction mechanisms that are often more difficult to characterize than interactions between conventional small-molecule drugs alone.

Although experimental studies and clinical trials remain the gold standard for identifying DDIs, comprehensive evaluation of all potential drug combinations is prohibitively expensive, time-consuming, and labor-intensive [8]. Consequently, computational methods have emerged as indispensable tools for prioritizing candidate interactions, accelerating drug discovery, and supporting clinical decision-making. Recent advances in artificial intelligence, particularly deep learning, graph neural networks, and hybrid learning frameworks, have significantly improved the prediction of drug–drug interactions, drug–target interactions, and adverse drug reactions by learning complex patterns directly from heterogeneous biomedical data [9]. A recent systematic review of machine learning and deep learning methods for anticancer drug interactions highlighted the rapid adoption of graph neural networks, Transformer-based models, and multimodal learning for DDI prediction, while identifying limited interpretability and poor generalization as major challenges that remain to be addressed [10]. Nevertheless, important challenges remain, including the limited availability of high-quality annotated datasets, severe class imbalance, increasing data heterogeneity, and the need for more accurate and interpretable predictive models [9].

Although considerable progress has been achieved in computational DDI prediction, several important challenges remain. Unlike small-molecule drugs, which have simple chemical structures and can be easily represented using SMILES, biotech drugs are large and complex molecules such as proteins and antibodies. Their biological properties, target-specific mechanisms, and complex pharmacokinetics make their interactions with small-molecule drugs more difficult to predict. In addition, the limited availability of biotech–small-molecule interaction data further increases the challenges of developing accurate computational prediction models [11]. Most existing studies have primarily focused on interactions between conventional small-molecule drugs, while comparatively little attention has been devoted to interactions involving biologic therapeutics despite their increasing clinical use [11–13]. Furthermore, many state-of-the-art approaches rely on predefined similarity measures, molecular interaction networks, or graph-based representations, which may limit their ability to generalize to previously unseen drug pairs and fully exploit heterogeneous molecular information [11, 14]. In addition, the majority of existing deep learning models prioritize predictive performance without providing sufficient interpretability, making it difficult to understand the molecular features underlying their predictions and limiting their applicability in biomedical research and clinical decision-making [9, 15].

Taken together, the existing literature indicates that an effective framework for predicting interactions between biotech and small-molecule drugs should satisfy several key requirements. First, it should integrate complementary molecular representations capable of capturing the diverse structural and biological characteristics of heterogeneous drug types. Second, it should effectively model both local and global dependencies to improve the characterization of complex interaction mechanisms. Third, it should accurately predict specific interaction types while maintaining robust generalization to previously unseen drug pairs. Finally, as predictive models become increasingly integrated into biomedical research and clinical practice, explainability has emerged as a critical requirement for improving model transparency, facilitating biological interpretation, and increasing users’ confidence in computational predictions [16, 17]. Addressing these challenges remains an important direction for the development of next-generation computational DDI prediction models.

To address the aforementioned challenges, we propose the Biotech–Small-Molecule Attention ResNet Transformer, called B-SMART-Former, an explainable deep learning framework for predicting interaction types between biotech and small-molecule drugs. The proposed framework integrates multimodal molecular representations by combining ChemBERTa embeddings and molecular fingerprints for small-molecule drugs with ProtBERT embeddings for biotech drugs. These complementary representations are processed through a hybrid neural architecture that combines Transformer-based self-attention with residual convolutional learning to capture both global contextual dependencies and local discriminative patterns. B-SMART-Former avoids using similarity-based features to improve its ability to generalize to previously unseen biotech and small-molecule drugs. Furthermore, Integrated Gradients is employed as a post-hoc explainability technique to identify the molecular features that contribute most significantly to the model’s predictions, thereby improving transparency and interpretability. The main contributions of this work are summarized as follows:

- Utilizing foundation model representations for both small-molecule and biotech drugs as input features to the proposed model.
- Eliminating similarity-based features to improve the generalization capability of the model, particularly for unseen drugs, without requiring feature recomputation.
- Integrating complementary protein and molecular language model representations within a unified Transformer-ResNet architecture.
- Proposing a novel architecture that integrates Transformer and residual convolutional networks to generate informative drug embeddings.
- Applying the post-hoc explainability method, Integrated Gradients, to identify the molecular features that most strongly influence the model’s predictions.
- Conducting comprehensive comparative and ablation studies to evaluate the effectiveness and robustness of B-SMART-Former.

The remainder of this paper is organized as follows. Section 3 reviews the related work. Section 4 presents the proposed methodology. Section 5 reports the experimental results, including the explainability analysis. Finally, Section 6 presents the discussion and conclusion, summarizes the main findings, discusses their implications, and outlines directions for future research.

## 3 Related work

This section reviews the literature related to computational drug–drug interaction prediction. We first discuss studies on drug–drug interaction (DDI) prediction involving small-molecule drugs, as they constitute the majority of existing research. We then review studies that investigate interactions between small-molecule and biotech drugs, which is the primary focus of this work. Finally, we summarize post-hoc explainability methods for deep learning models.

### 3.1 Small-molecule DDI

Small-molecule drug interaction prediction has been the primary focus of computational DDI research during the past decade. According to the recent systematic review by Behrooznia and Hooshmand [9], computational approaches for drug interaction prediction can generally be categorized into conventional machine learning, deep learning, graph learning, and hybrid methods. Recent studies have increasingly shifted toward graph neural networks and Transformer-based architectures because of their superior ability to model complex molecular relationships.

Traditional computational approaches for drug–drug interaction (DDI) prediction primarily relied on conventional machine learning algorithms, including Support Vector Machines (SVMs), Random Forests (RFs), Logistic Regression (LR), and gradient boosting methods. These approaches typically employed handcrafted features derived from molecular fingerprints, chemical structures, drug targets, biological pathways, and therapeutic information to predict potential interactions [5].

Although these methods demonstrated promising predictive performance, they relied heavily on manually engineered features and exhibited limited capability to capture complex nonlinear relationships from heterogeneous molecular and biological data. Consequently, recent research has shifted toward deep learning techniques, which automatically learn hierarchical feature representations and provide improved generalization for DDI prediction [18].

The emergence of deep learning has substantially advanced DDI prediction by enabling end-to-end representation learning from diverse molecular and biological data sources. Architectures such as multilayer perceptrons (MLPs), convolutional neural networks (CNNs), recurrent neural networks (RNNs), and attention-based models have consistently outperformed conventional machine learning approaches by learning discriminative features directly from raw inputs [15].

Despite these advances, existing deep learning methods still face several challenges. Many approaches focus on local feature extraction or rely on a single modality of drug representation, limiting their ability to capture complex dependencies and fully exploit heterogeneous information. These limitations have encouraged the development of more expressive graph neural network (GNN) and Transformer-based architectures [14].

Graph neural networks have become one of the dominant paradigms for DDI prediction by naturally representing drugs and their interactions as graph-structured data. Li *et al*. proposed MDG-DDI, which constructs a multi-feature drug graph by integrating biochemical properties with interaction network information and employs graph neural networks together with a self-attention module to learn informative drug representations [19]. Gong *et al*. introduced MDI-DDI, a multi-dimensional interaction graph neural network that utilizes GraphSAGE convolutions and contrastive learning to capture complementary structural and semantic relationships between drugs [20]. Nishio *et al*. further demonstrated the effectiveness of graph-based representation learning by integrating DrugBank molecular information within a GNN framework to jointly predict drug–drug and drug–food interactions [21]. Nevertheless, the message-passing mechanism of GNNs is primarily based on local neighborhood aggregation, which may limit their ability to capture long-range dependencies between heterogeneous drug features [22].

To overcome these limitations, recent studies have increasingly adopted attention- and Transformer-based architectures to model global feature interactions. Liu *et al*. proposed DANN-DDI, which employs a deep attention neural network to adaptively weight multiple drug similarity features before interaction prediction [23]. Su and Qian introduced DDI-Transform, a Transformer-based framework that integrates drug structural information with drug-binding protein features using self-attention to learn comprehensive representations for DDI event prediction [24]. More recently, Tamir and Yuan proposed KITE-DDI, which combines SMILES sequences with biomedical knowledge graph information within a Transformer architecture, enabling the joint learning of molecular characteristics and semantic relationships for more accurate DDI prediction [25].

Xia *et al*. [26] proposed MFDL-DDI, a multimodal deep learning framework that integrates heterogeneous drug information through a multimodal feature fusion strategy for DDI prediction. Specifically, the framework combines molecular structures, biological attributes, and drug similarity information to learn complementary representations from multiple data modalities. Experimental results demonstrated that multimodal information fusion improves the predictive performance of DDI prediction.

Zou *et al*. [27] proposed MS-DDI, a multi-scale representation learning framework for drug–drug interaction prediction. The model captures drug characteristics at multiple representation scales and integrates them to learn comprehensive interaction patterns, thereby improving the discriminative capability of the learned drug representations. While these methods have achieved promising performance, they primarily focus on interactions between small-molecule drugs and provide limited model interpretability.

### 3.2 Biotech–small-molecule DDI

Compared with small-molecule DDIs, studies investigating interactions between biotech and small-molecule drugs remain relatively limited. Huang *et al*. [12] proposed a three-stage framework that extracts structural and interaction-related features from both drug types, generates feature vectors using one-hot encoding, Jaccard similarity, and principal component analysis (PCA), and subsequently employs a deep learning model for interaction prediction. However, Ru *et al*. [13] later demonstrated that the one-hot encoding and Jaccard similarity calculations were implemented incorrectly, resulting in PCA being applied to inaccurate feature representations. Furthermore, the reported experiments addressed only binary interaction prediction rather than the more challenging task of predicting specific interaction types.

To address these shortcomings, Ru *et al*. [13] proposed a graph-based framework built upon a two-level graph representation. The first level models drug–drug and drug–target relationships, whereas the second captures structural information for individual small-molecule drugs. Graph convolutional networks were employed to learn node embeddings, which were subsequently fed into a multilayer perceptron for interaction type prediction. However, despite targeting biotech–small-molecule interactions, the experimental evaluation primarily emphasized small-molecule DDIs, providing only limited validation of the intended prediction task.

More recently, Nasiri *et al*. proposed BSI-Net, a graph-based deep learning framework that integrates structural representations of biotech and small-molecule drugs with similarity information to predict interaction events [11]. By exploiting similarity matrices, BSI-Net effectively models relationships between heterogeneous drug types and achieves competitive predictive performance [28]. Nevertheless, its dependence on predefined similarity measures may limit scalability and reduce its ability to generalize to previously unseen drug pairs.

### 3.3 Explainable artificial intelligence for DDI prediction

Although predictive performance has improved considerably, model interpretability has only recently become an active research topic in computational DDI prediction. The importance of model interpretability in computational DDI prediction has received increasing attention in recent years. A recent systematic review by Liu *et al*. [29] highlighted that, although graph neural networks and Transformer-based models have significantly improved predictive performance, most existing approaches remain black-box models, making it difficult to understand the molecular mechanisms underlying their predictions. The review also highlights the growing importance of model interpretability and explainability for developing trustworthy DDI prediction systems.

Among the existing explainable DDI models, Kim and Nam [30] proposed DeSIDE-DDI, an interpretable deep learning framework that predicts drug–drug interactions from drug-induced gene expression profiles. The model employs a gating mechanism that dynamically assigns importance to gene expression features, enabling the identification of genes that contribute most to each prediction. The resulting importance scores are subsequently analyzed to highlight the gene expression signatures responsible for the predicted interaction, thereby providing biologically meaningful explanations alongside accurate predictions.

Wang *et al*. [31] proposed KnowDDI, an interpretable graph neural network framework that combines biomedical knowledge graphs with knowledge subgraph learning for DDI prediction. The model first learns drug representations by propagating information over a biomedical knowledge graph using graph neural networks and subsequently extracts interaction-specific knowledge subgraphs to explain individual predictions. Instead of relying on feature attribution, KnowDDI explains its decisions by identifying the biomedical entities and relationships that contribute most to the predicted interaction.

Kha *et al*. [32] developed a machine learning framework for predicting DDIs associated with oral diabetes medications. To improve model interpretability, they employed SHAP to quantify the contribution of individual molecular features, enabling both local and global explanations of the model’s predictions.

Existing explainable DDI studies primarily rely on attention mechanisms or knowledge graph reasoning to improve model interpretability. Compared with these explanation strategies, the application of gradient-based attribution methods, such as Integrated Gradients, remains relatively limited in DDI prediction, particularly for interaction type prediction involving biotech and small-molecule drugs.

Table 1 summarizes the representative studies reviewed in this section, highlighting their methodological characteristics, explainability strategies, and primary limitations.

**Table 1.** Summary of representative studies on computational drug–drug interaction prediction.

| Method | Year | Drug type | Main methodology | Explainability |
| --- | --- | --- | --- | --- |
| DANN-DDI [23] | 2022 | SM-SM | Deep attention network with multiple drug similarity features. | No |
| DeSIDE-DDI [30] | 2022 | SM-SM | Gene expression-based prediction with attention gating. | Attention-based |
| DDI-Transform [24] | 2024 | SM-SM | Transformer integrating structural and protein-binding features. | No |
| KnowDDI [31] | 2024 | SM-SM | Knowledge graph learning with interpretable subgraph reasoning. | Knowledge graph reasoning |
| Kha *et al.* [32] | 2024 | SM-SM | Machine learning with SHAP-based feature attribution. | SHAP |
| KITE-DDI [25] | 2025 | SM-SM | Transformer combining SMILES and biomedical knowledge graphs. | No |
| MDG-DDI [19] | 2025 | SM-SM | Multi-feature graph neural network with self-attention. | No |
| MDI-DDI [20] | 2025 | SM-SM | GraphSAGE and contrastive learning for graph representation learning. | No |
| MFDL-DDI [26] | 2026 | SM-SM | Multimodal fusion of molecular, biological, and similarity features. | No |
| MS-DDI [27] | 2026 | SM-SM | Multi-scale representation learning for drug interaction prediction. | No |
| Multi-SBI [12] | 2022 | Biotech-SM | One-hot encoding, Jaccard similarity, PCA, and deep learning. | No |
| CB-TIP [13] | 2023 | Biotech-SM | Two-level graph representation with GCN and MLP classifier. | No |
| BSI-Net [11] | 2025 | Biotech-SM | Graph-based learning with structural and similarity representations. | No |

Collectively, the reviewed studies demonstrate that, despite substantial progress in computational DDI prediction, several important challenges remain. Existing studies have predominantly focused on interactions between small-molecule drugs, while biotech–small-molecule DDIs remain comparatively underexplored. Furthermore, many current approaches either rely heavily on handcrafted similarity measures or provide limited interpretability of their predictions. These limitations highlight the need for models capable of learning discriminative multimodal representations, capturing both local and global dependencies among heterogeneous drug features, and providing transparent, biologically meaningful explanations for their predictions.

## 4 Materials and methods

### 4.1 Dataset

The dataset used in this study was derived from previous work [11], which was originally curated from the DrugBank database [8]. In this work, we focus on biotech–small-molecule drug pairs. The dataset consists of 196 approved biotech drugs and 2,148 approved small-molecule drugs. The final prediction task includes a total of 87,752 biotech–small-molecule drug pairs, comprising 43,876 positive interaction pairs and 43,876 negative pairs representing the absence of a known interaction. The negative interaction pairs were generated by randomly sampling biotech–small-molecule drug pairs that were not reported as interacting in DrugBank. To obtain a balanced dataset, the number of negative samples was matched to the number of positive interaction pairs while ensuring that no sampled negative pair overlapped with the known positive interactions. Each sample consists of one biotech drug, one small-molecule drug, and a corresponding interaction label. The interaction prediction problem is formulated as a multi-class classification task with 32 labels, consisting of one negative class (Label 0) and 31 positive interaction types (Labels 1–31). The 31 positive interaction labels were selected from a larger set of interaction types. The selected labels account for more than 97% of all available labeled interactions. Interaction labels with fewer than 50 occurrences were excluded because they do not provide sufficient samples for reliable model training and evaluation. Furthermore, most interaction labels are bidirectional. Although DrugBank contains 62 textual interaction descriptions, most describe symmetric interactions. Therefore, each bidirectional interaction pair is represented by a single interaction label in this study. For example, if Drug 1 has interaction type *x* with Drug 2, then Drug 2 is also considered to have interaction type *x* with Drug 1. The negative class represents the absence of a known interaction between a biotech and a small-molecule drug, whereas the positive labels correspond to diverse clinically relevant interaction events extracted from DrugBank. These interaction types encompass a broad range of pharmacokinetic and pharmacodynamic effects, including alterations in drug metabolism, excretion, serum concentration, therapeutic efficacy, anticoagulant activity, and toxicity. Table 2 summarizes the dataset statistics, while representative interaction labels are presented in Table 3. For each small-molecule drug *i*, where *i* ∈ {1, …, *n*} and *n* denotes the total number of small-molecule drugs, the corresponding SMILES representation, denoted by SM_*i*_, is used as the input feature. Similarly, for each biotech drug *j*, where *j* ∈ {1, …, *m*} and *m* denotes the total number of biotech drugs, the corresponding amino acid sequence, denoted by BT_*j*_, is used as the input feature. The corresponding interaction label is denoted by *y*_*ij*_ ∈ {0, 1, …, 31}, where 0 represents the negative class (no known interaction) and labels 1–31 denote the positive interaction types.

**Table 2.** Dataset statistics.

| Entity | Frequency |
| --- | --- |
| Biotech drugs | 196 |
| Small-molecule drugs | 2,148 |
| Positive interaction pairs | 43,876 |
| Negative interaction pairs | 43,876 |
| Total interaction pairs | 87,752 |
| Positive interaction labels | 31 |
| Negative interaction label | 1 |
| Total labels | 32 |

**Table 3.** Representative interaction labels used in the dataset.

| Label | Description |
| --- | --- |
| 0 | Negative or unknown interaction (no known interaction between the biotech and small-molecule drug). |
| 1 | The metabolism of drug2 can be increased when combined with drug1 (bidirectional case). |
| 2 | The risk or severity of adverse effects can be increased when drug2 is combined with drug1 (bidirectional case). |
| ⋮ | ⋮ |
| 31 | drug2 may decrease the anticoagulant activities of drug1 (bidirectional case). |

### 4.2 Feature representation

For each drug pair (*i, j*), where *i* and *j* denote the indices of the small-molecule and biotech drugs, respectively, the proposed B-SMART-Former constructs a unified feature vector, denoted by **f**_*ij*_, by combining the representations of the two drugs. This multimodal representation integrates complementary chemical and biological information to provide a comprehensive characterization of each biotech–small-molecule drug pair.

#### Small-molecule representation

Small-molecule drugs were represented using a hybrid feature vector composed of ChemBERTa embeddings and circular molecular (Morgan) fingerprints. Specifically, the SMILES string of each small molecule was encoded using the pretrained `seyonec/ChemBERTa_zinc250k_v2_40k` model, a Transformer-based molecular language model [33, 34] trained on large-scale SMILES corpora.

Let *S* = {*s*_1_, *s*_2_, …, *s*_*ℓ*_} denote the tokenized SMILES sequence of a small-molecule drug, where *ℓ* denotes the sequence length. Each token is first mapped into a continuous embedding space by combining its token embedding with the corresponding positional embedding:

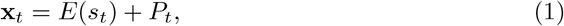

where *E*(·) denotes the token embedding function and *P*_*t*_ represents the positional embedding of the *t*-th token.

The resulting embedding sequence is processed by the stacked Transformer encoder layers [33] of ChemBERTa to learn contextual representations of the SMILES tokens:

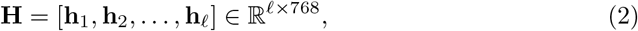

where **h**_*t*_ denotes the contextual representation of the *t*-th SMILES token. The final ChemBERTa embedding is obtained by applying mean pooling over all hidden states:

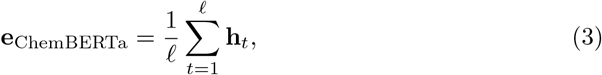

where **e**_ChemBEℝTa_ ℝ^768^ denotes the final molecular embedding.

To complement the contextual molecular representation learned by ChemBERTa, circular molecular (Morgan) fingerprints [35] were generated using a radius of 2 and a fingerprint size of 512 bits. Morgan fingerprints encode circular atomic neighborhoods around each atom, thereby capturing local molecular substructures and providing a compact representation of molecular topology [36].

Let *G* = (*V, E*) denote the molecular graph of a small-molecule drug, where *V* and *E* represent the sets of atoms and chemical bonds, respectively. The corresponding Morgan fingerprint is computed as

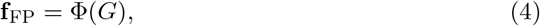

where Φ(·) denotes the Morgan fingerprint generation procedure with radius *r* = 2 and fingerprint dimensionality *d* = 512, and **f**_FP_ ∈ {0, 1}^512^ is the resulting binary fingerprint vector.

The final representation of the *i*-th small-molecule drug is obtained by concatenating the ChemBERTa embedding and the Morgan fingerprint:

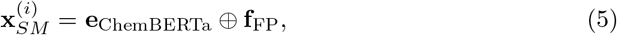

where ⊕ denotes vector concatenation and 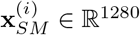.

#### Biotech drug representation

Biotech drugs were represented using fixed-length embeddings extracted from their amino acid sequences with the pretrained ProtBERT model [37]. ProtBERT is a Transformer-based protein language model trained to learn contextual representations of protein sequences, enabling it to capture both local and long-range dependencies among amino acid residues.

Let 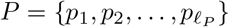 denote the amino acid sequence of the *j*-th biotech drug, where *ℓ*_*P*_ denotes the protein sequence length. Similar to ChemBERTa, the sequence is processed by the pretrained ProtBERT encoder to generate contextual representations for each amino acid residue. The resulting hidden-state matrix is given by

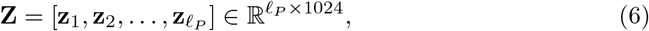

where **z**_*k*_ ∈ ℝ^1024^ denotes the contextual representation of the *k*-th amino acid residue.

To obtain a fixed-length representation suitable for downstream classification, mean pooling is applied over the residue representations:

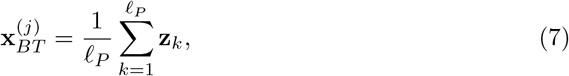

where 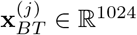 denotes the final embedding of the *j*-th biotech drug.

#### Drug-pair representation

The final multimodal representation of the interaction between the *i*-th small-molecule drug and the *j*-th biotech drug is obtained by concatenating their respective feature vectors:

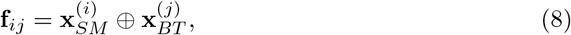

where **f**_*ij*_ ∈ ℝ^2304^ denotes the final feature vector representing the biotech–small-molecule drug pair.

Before model training, the concatenated feature vectors were standardized using the `StandardScaler` algorithm. Within each fold of the stratified 10-fold cross-validation procedure, the scaler was fitted exclusively on the training data and subsequently applied to the corresponding validation and test sets. This preprocessing strategy prevents information leakage and ensures an unbiased evaluation of the proposed framework [38].

### 4.3 Proposed model

We propose B-SMART-Former, a deep learning model designed to predict interaction types between biotech and small-molecule drugs. The model integrates dense projection layers, a Transformer-based module, residual convolutional blocks, and a final multi-layer perceptron classifier to learn complementary global and local feature representations. This section describes the details of the framework. Figure 1 illustrates the general structure of B-SMART-Former.

**Fig 1.**
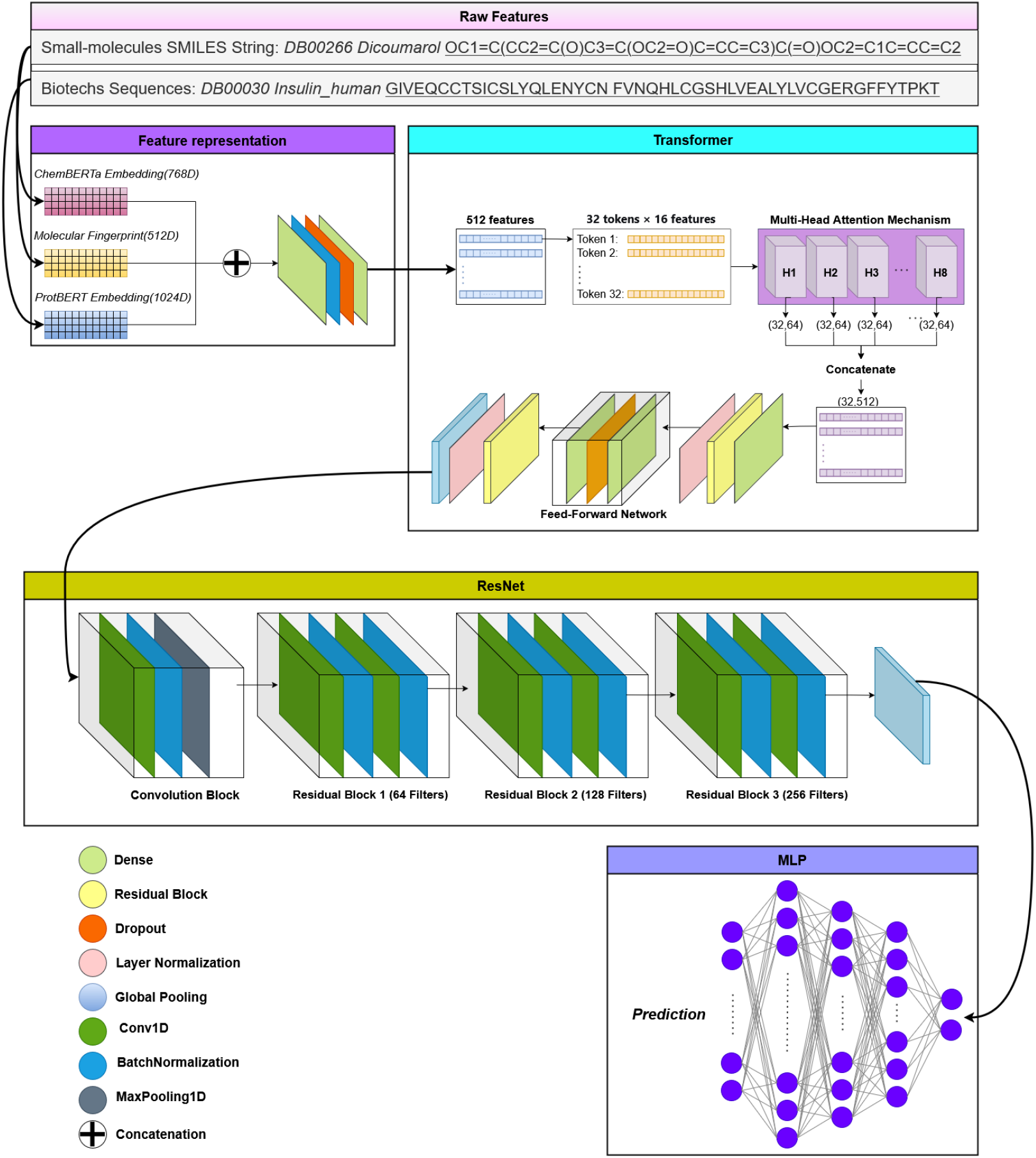
Overview of the proposed B-SMART-Former framework for biotech–small-molecule drug–drug interaction prediction. The model takes the SMILES representation of a small-molecule drug and the amino acid sequence of a biotech drug as inputs. The SMILES representation is encoded using ChemBERTa and Morgan fingerprints, while the biotech sequence is encoded using ProtBERT. The resulting feature representations are concatenated and passed through a feature projection module to reduce the input dimensionality. The projected features are then processed by a Transformer encoder consisting of a multi-head self-attention layer and a feed-forward network, followed by a residual convolutional (ResNet) module to generate informative joint embeddings. Finally, a multi-layer perceptron (MLP) predicts the interaction type for the biotech–small-molecule drug pair.

The model takes the concatenated drug-pair feature vector as input and first projects it into a lower-dimensional space using dense layers. The representation is then reshaped into a sequential format and processed by a Transformer module to capture global dependencies among features. The resulting representation is further refined using residual convolutional blocks to extract local patterns. Finally, a multi-layer perceptron produces the predicted interaction class.

In the following subsections, each component of the proposed architecture is described in detail.

#### 4.3.1 Input projection layers

The input to B-SMART-Former is the concatenated feature vector **f**_*ij*_ ∈ ℝ^2304^, representing the feature embeddings of a biotech–small-molecule drug pair and constructed according to Eq. (8) in the feature representation stage. Since this vector integrates heterogeneous molecular and biological information extracted from different modalities, it is first projected into a lower-dimensional latent space before sequence modeling. This projection enables the network to learn nonlinear interactions among the input features while producing a compact and discriminative representation for subsequent processing.

To achieve this, two fully connected projection layers are employed. Each dense layer uses the Swish activation function [39], followed by batch normalization and dropout to improve optimization stability, accelerate convergence, and reduce overfitting. For the proposed input representation of 2,304 dimensions, the projection layers reduce the feature dimensionality from 2,304 to 1,024 and subsequently to 512 dimensions.

The projection process can be formulated as

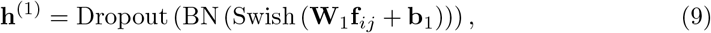

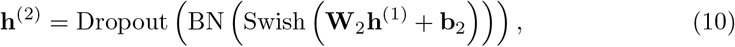

where **W**_1_ ∈ ℝ^1024*×*2304^ and **W**_2_ ∈ ℝ^512*×*1024^ denote the trainable weight matrices, while **b**_1_ and **b**_2_ represent the corresponding bias vectors. The operator BN(·) denotes batch normalization, Dropout(·) represents the dropout operation, and Swish(·) is the Swish activation function.

The resulting latent representation **h**^(2)^ ∈ ℝ^512^ serves as the input to the Transformer module. Before being processed by the Transformer encoder, this representation is reshaped into a sequence, enabling the model to capture contextual dependencies among latent feature groups through self-attention.

#### 4.3.2 Transformer module

The Transformer module is employed to model global dependencies among the latent feature representations using the self-attention mechanism [33]. Unlike convolutional layers, which capture local patterns within a limited receptive field, self-attention enables the model to learn relationships between all feature groups simultaneously, thereby capturing long-range interactions among heterogeneous drug features.

Following the input projection stage, the latent representation **h**^(2)^ ∈ ℝ^512^ is reshaped into a sequence of feature tokens before being processed by the Transformer encoder. To preserve the ordering of these tokens, learnable positional embeddings are added to the input sequence. Let

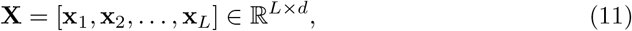

denote the resulting sequence, where *L* is the sequence length and *d* is the embedding dimension of each token.

For each Transformer encoder block, the query, key, and value matrices are obtained through linear projections,

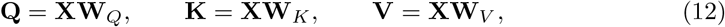

where **W**_*Q*_, **W**_*K*_, and **W**_*V*_ are trainable projection matrices.

The scaled dot-product self-attention is computed as

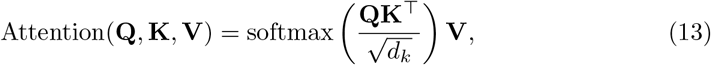

where *d*_*k*_ denotes the dimensionality of the key vectors.

To capture information from multiple representation subspaces, multi-head self-attention is employed:

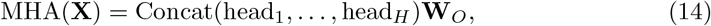

where

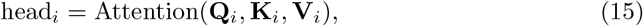

and *H* denotes the number of attention heads.

Each Transformer encoder block further incorporates a position-wise feed-forward network (FFN), residual connections, and layer normalization to improve optimization stability and facilitate gradient propagation. In the proposed architecture, two Transformer encoder blocks are employed, using eight and four attention heads, respectively. The resulting contextual feature sequence is finally aggregated by a Global Average Pooling layer to produce a compact global representation for the subsequent residual convolutional module.

#### 4.3.3 Residual convolutional module

Although the Transformer module effectively captures global contextual dependencies through self-attention, it is less specialized in modeling local feature interactions. To complement the Transformer representation, B-SMART-Former incorporates a residual convolutional module that extracts local and hierarchical feature patterns from the contextual sequence.

Let **T** ∈ ℝ^*L*×*d*^ denote the output of the Transformer encoder, where *L* is the sequence length and *d* is the feature dimension. The input sequence is processed using one-dimensional convolutional layers, which capture local dependencies by applying learnable filters over neighboring feature tokens [40]. The convolutional operation can be expressed as

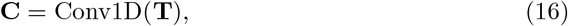

where **C** denotes the resulting local feature representation.

To facilitate efficient optimization and preserve information learned by the Transformer module, residual skip connections are introduced between the convolutional layers [41]. The output of a residual block is computed as

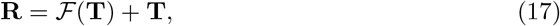

where *F* (·) represents the nonlinear transformation performed by the stacked convolutional layers, including the Swish activation function and batch normalization. The residual connection enables the network to learn residual mappings instead of complete feature transformations, thereby improving gradient propagation and reducing the risk of performance degradation in deeper architectures.

Finally, the output of the residual convolutional module is aggregated using Global Average Pooling to obtain a compact feature representation, which is subsequently forwarded to the classification module.

#### 4.3.4 Classification head

The global representations learned by the Transformer and residual convolutional modules capture complementary contextual and local feature information. To exploit both representations jointly, their pooled outputs are concatenated to form a unified feature vector,

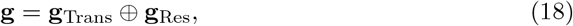

where **g**_Trans_ and **g**_ℝes_ denote the Global Average Pooling outputs of the Transformer and residual convolutional branches, respectively.

The fused representation is subsequently forwarded to a multi-layer perceptron (MLP) classifier consisting of three fully connected layers. Each hidden layer employs the Swish activation function [39], followed by batch normalization and dropout to improve optimization stability, enhance generalization, and reduce overfitting. This hierarchical transformation enables the classifier to learn increasingly discriminative representations for interaction prediction.

The hidden representations are computed as

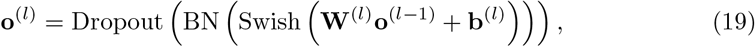

where **o**^(0)^ = **g**, and **W**^(*l*)^ and **b**^(*l*)^ denote the trainable weight matrix and bias vector of the *l*-th hidden layer, respectively.

The final output layer employs the Softmax activation function to estimate the probability distribution over the 32 interaction classes:

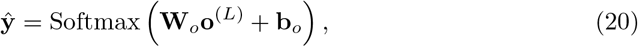

where ***ŷ*** ∈ ℝ^32^ denotes the predicted probability distribution over all interaction classes, and *L* represents the final hidden layer of the MLP.

#### 4.3.5 Model training

The proposed B-SMART-Former was trained as a multi-class classification model using the sparse categorical cross-entropy loss function,

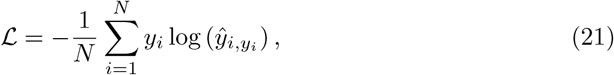

where *N* denotes the number of training samples, *y*_*i*_ is the ground-truth interaction label of the *i*-th sample, and 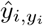 is the predicted probability assigned to the corresponding class.

Model parameters were optimized using the Adam optimizer [42] with a cosine decay restart learning-rate schedule [43]. This adaptive optimization strategy accelerates convergence while periodically restarting the learning rate to improve exploration of the optimization landscape.

To mitigate the effect of class imbalance, class weights were computed from the training data of each fold using a balanced weighting strategy and incorporated into the loss function during optimization. Furthermore, early stopping based on the validation loss was employed to prevent overfitting by restoring the model parameters corresponding to the best-performing epoch.

### 4.4 Training procedure

The proposed framework was evaluated using stratified 10-fold cross-validation to obtain a robust and unbiased estimate of its predictive performance [44]. Stratification preserved the original class distribution across all folds, ensuring representative training and testing subsets for every interaction class.

For each fold, the dataset was divided into training and testing subsets. Subsequently, 10% of the training samples were randomly selected as a validation set for model selection and performance monitoring during training.

To prevent information leakage, feature standardization was performed independently within each fold. Specifically, a `StandardScaler` was fitted exclusively on the training data and then applied to the corresponding validation and test sets.

The network was trained for a maximum of 30 epochs using a batch size of 16. During training, model checkpointing was employed to preserve the network parameters corresponding to the lowest validation loss. After optimization, the best-performing model from each fold was evaluated on the held-out test set. The final performance metrics reported in this study correspond to the average results obtained across all ten cross-validation folds.

Algorithm 1 outlines the complete training and evaluation workflow of the proposed B-SMART-Former framework.

Lines 1–2 define the algorithm inputs, including the precomputed drug-pair feature matrix, the corresponding interaction labels, and the number of cross-validation folds, while the output consists of the predicted interaction labels.

Lines 3–7 describe the data preparation stage performed within each fold of the stratified cross-validation procedure. The dataset is divided into training and testing subsets, a validation subset is created from the training data, feature standardization is performed using a StandardScaler fitted only on the training set, and class weights are computed to alleviate the effect of class imbalance.

Lines 8–11 summarize the forward propagation through the proposed B-SMART-Former architecture. The standardized input features are first projected into a latent representation, followed by the Transformer module for global dependency modeling, the residual convolutional module for local feature extraction, and finally the MLP classifier for interaction prediction.

Lines 12–14 describe the optimization process. The network is trained using the sparse categorical cross-entropy loss function and optimized with the Adam optimizer using a cosine decay restart learning-rate schedule. Early stopping together with model checkpointing is employed to prevent overfitting and preserve the best-performing model.

Finally, Lines 15–18 evaluate the optimized model on the test subset of the current fold, store the predicted labels and evaluation metrics, and compute the average predictive performance across all cross-validation folds.

#### Algorithm 1 Training pipeline of the proposed B-SMART-Former

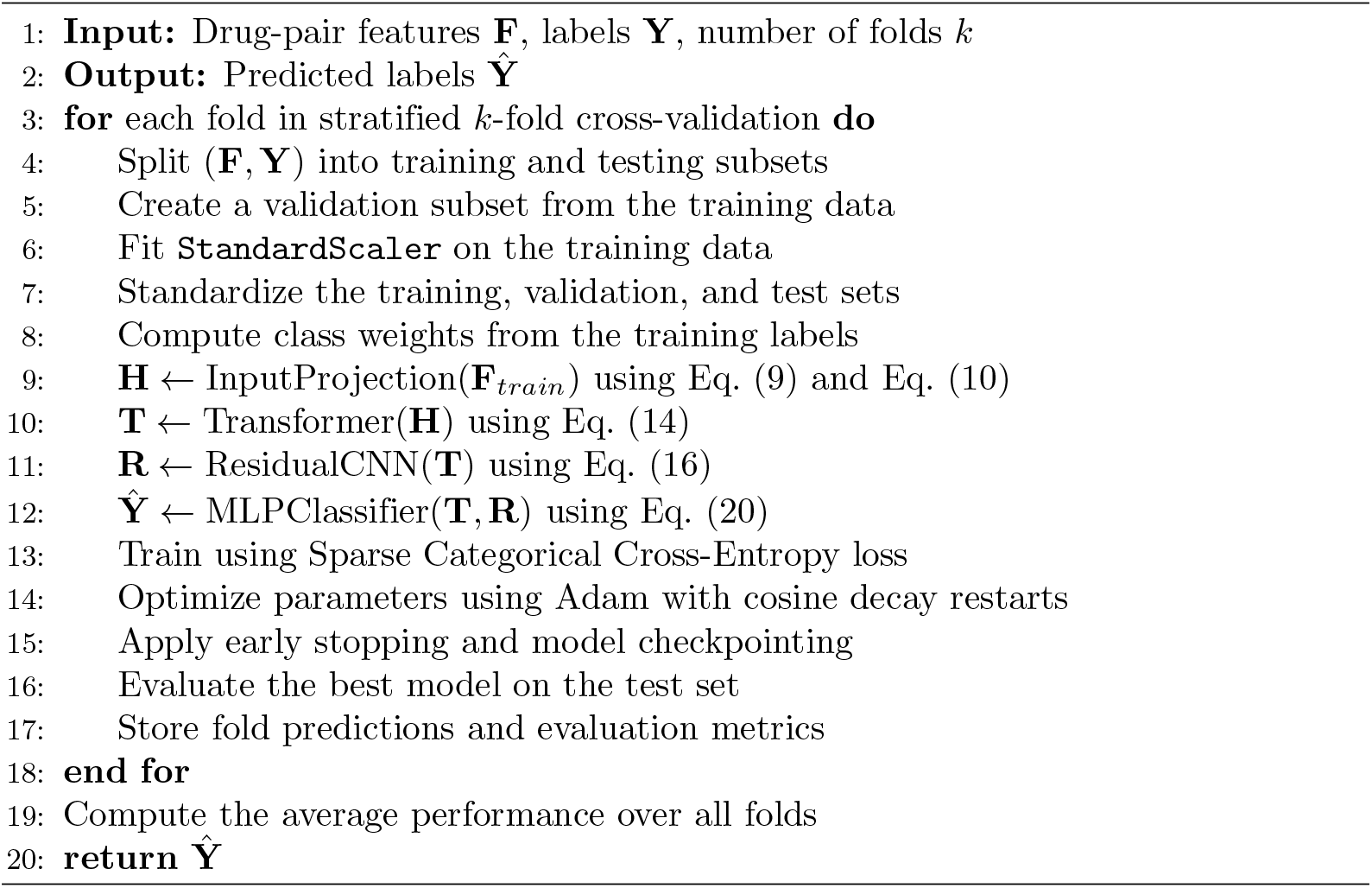

### 4.5 Explainability analysis

Deep learning models have achieved remarkable success in computational drug discovery by providing highly accurate predictions for various tasks. However, their complex architectures often behave as black-box models, making it difficult to understand the rationale behind their predictions. This lack of transparency limits their applicability in safety-critical applications such as drug–drug interaction prediction, where reliable and interpretable decision-making is essential. Explainable artificial intelligence (XAI) addresses this challenge by identifying the input features that contribute most to a model’s predictions and has been increasingly adopted in computational drug discovery to improve model transparency and trustworthiness [45].

XAI approaches are generally categorized into intrinsic and post-hoc methods. Intrinsic approaches employ inherently interpretable models, such as linear models and decision trees, whereas post-hoc approaches generate explanations for previously trained models without modifying their underlying architectures. Representative post-hoc techniques include SHapley Additive exPlanations (SHAP) [46], Local Interpretable Model-Agnostic Explanations (LIME) [17], and gradient-based attribution methods. These techniques provide meaningful explanations while preserving the predictive capability and flexibility of deep learning models.

In this study, a post-hoc explainability framework based on Integrated Gradients (IG) [16] is employed to quantify the contribution of individual input features to the model predictions. Integrated Gradients computes feature attribution scores by interpolating between a baseline input, which is set to a zero vector, and the actual input sample. Along this interpolation path, the gradients of the model output with respect to the input features are accumulated and scaled by the difference between the input and the baseline. Consequently, each input feature is assigned an attribution score that reflects its contribution to the final prediction.

Formally, the Integrated Gradients attribution for the *i*-th input feature is defined as

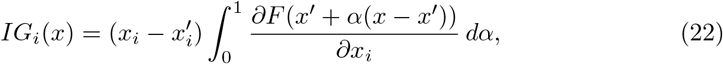

where *x* denotes the input sample, *x*^*′*^ represents the baseline input, *F* (·) is the prediction function of the trained B-SMART-Former model, and *α* ∈ [0, 1] is the interpolation coefficient. In practice, the integral is approximated using a finite number of interpolation steps.

To facilitate interpretation at the modality level, the computed attribution scores were grouped according to the three feature components that constitute the input representation, namely ChemBERTa embeddings, Morgan fingerprint features, and ProtBERT embeddings. For each explained sample, the absolute attribution values within each feature group were summed and subsequently normalized to obtain their relative percentage contributions. This aggregation provides an intuitive estimate of the contribution of each feature modality to the final prediction.

To further investigate the molecular fingerprint component, the most influential Morgan fingerprint bits were identified based on their attribution scores. Local explanations were generated by selecting the highest-contributing fingerprint bits for each individual prediction, whereas global explanations were obtained by aggregating fingerprint attributions across all explained samples. Whenever possible, the identified fingerprint bits were mapped back to their corresponding molecular substructures using RDKit [47], enabling the generation of molecular highlight maps. These visualizations provide chemically interpretable explanations by revealing the molecular substructures that contribute most to the predicted drug–drug interactions. Figure 2 illustrates the overall workflow of the Integrated Gradients method.

**Fig 2.**
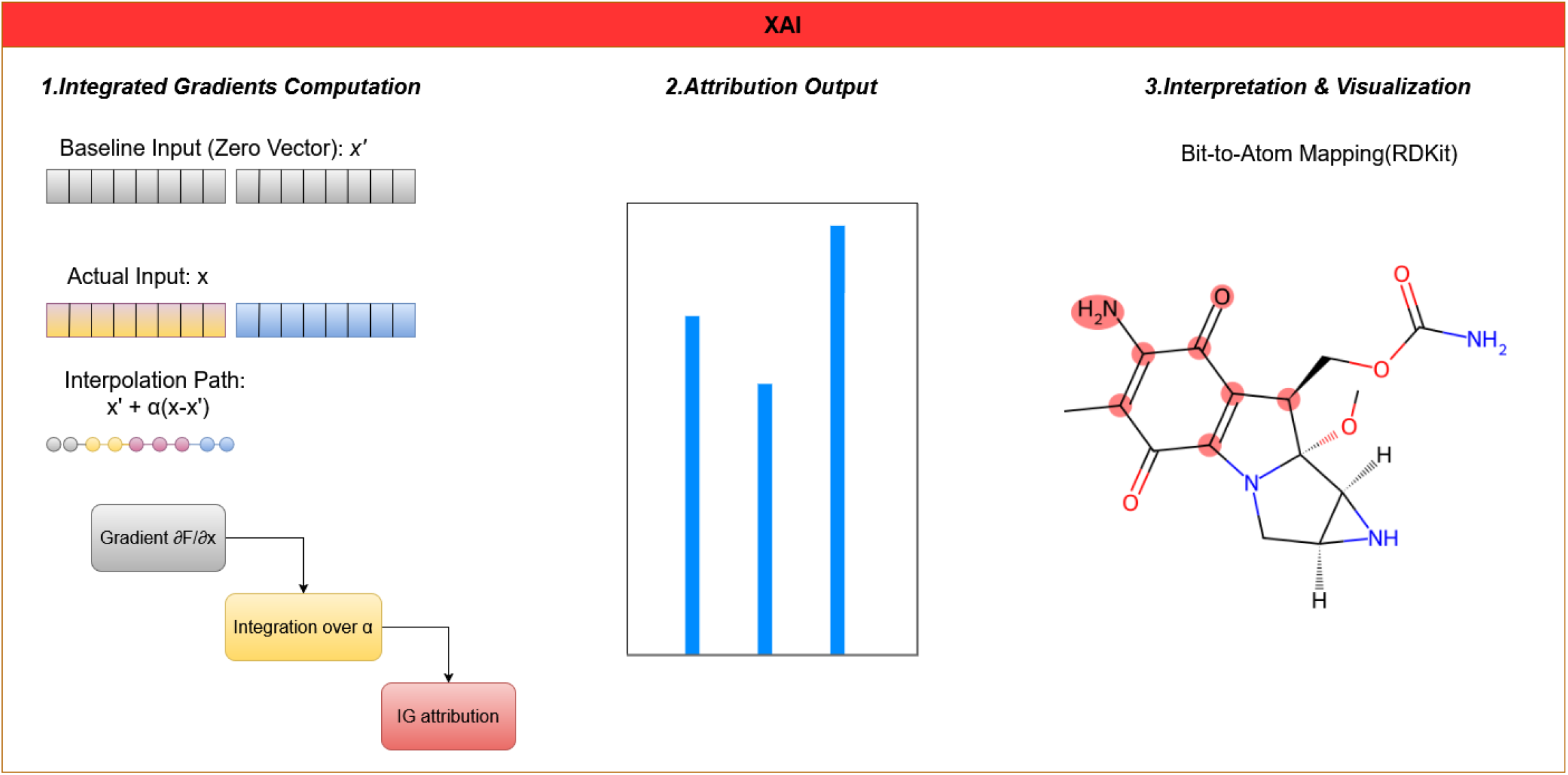
Overview of the Integrated Gradients-based explainability framework employed in B-SMART-Former. Starting from a zero-vector baseline, Integrated Gradients computes feature attribution scores by accumulating the gradients of the model output along the interpolation path to the original input. The resulting attributions quantify the contribution of individual input features to the predicted interaction. The most influential Morgan fingerprint bits are subsequently mapped to their corresponding molecular substructures using RDKit, enabling chemically interpretable visualization of the model’s predictions.

Algorithm 2 summarizes the post-hoc explainability workflow adopted in this study.

Lines 1–3 initialize the trained B-SMART-Former model and define the selected test samples used for explanation.

Lines 4–10 describe the local explanation process. For each test sample, a zero-valued baseline is generated and Integrated Gradients is applied to compute feature attribution scores. The resulting attributions are subsequently aggregated according to the three feature modalities, namely ChemBERTa, Morgan fingerprints, and ProtBERT, and normalized to obtain their relative contributions.

Finally, Lines 11–13 summarize the global explanation stage. Attribution scores are aggregated across all explained samples to estimate global feature importance. The most influential Morgan fingerprint bits are then identified and mapped back to molecular substructures using RDKit, providing chemically interpretable visual explanations of the model predictions.

#### Algorithm 2 Integrated Gradients-based explainability pipeline

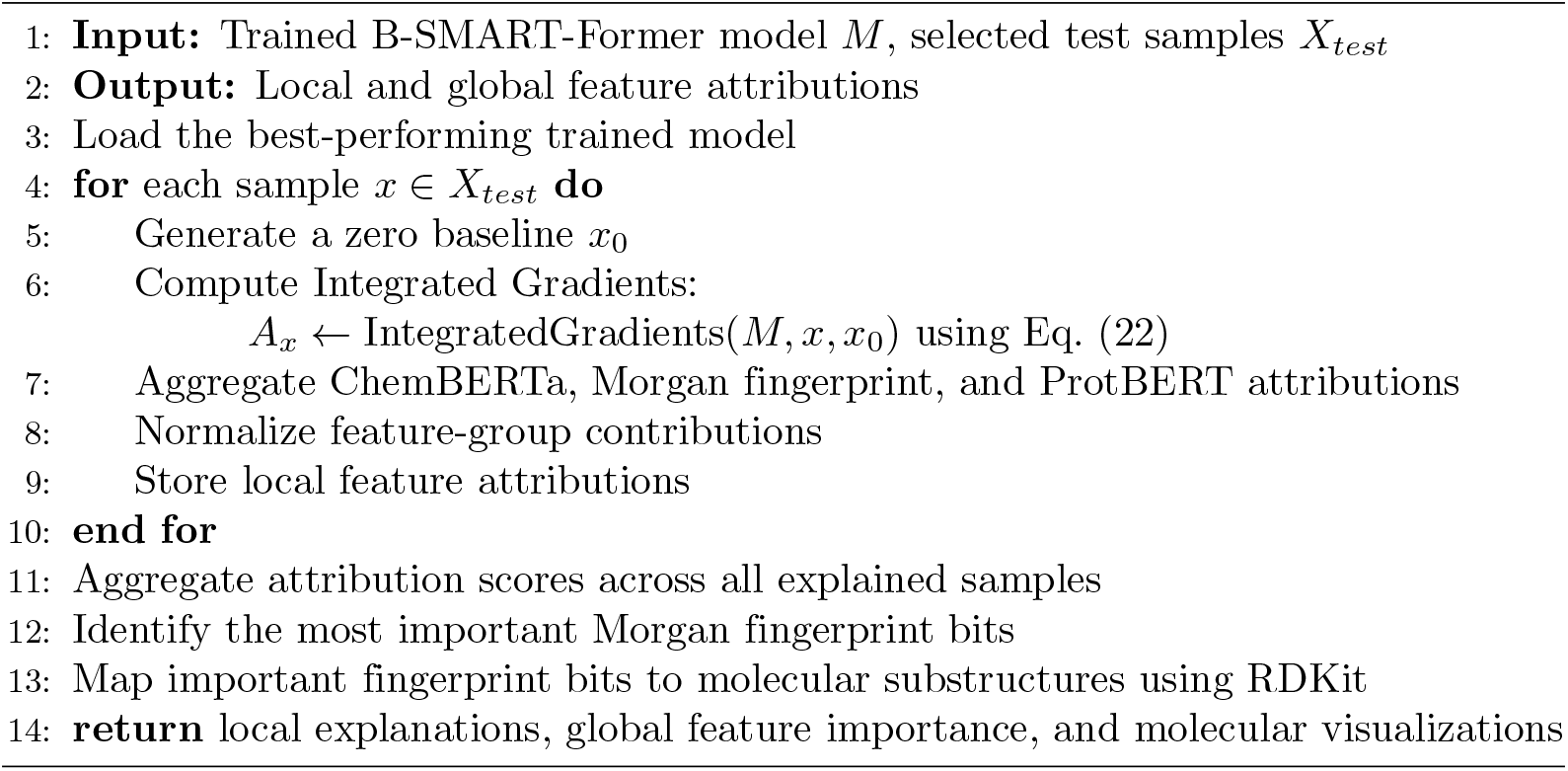

## 5. Results

All experiments were implemented in Python using the TensorFlow deep learning framework together with the Keras API for model development and training. Numerical computations were performed using NumPy, while the Scikit-learn library was utilized for data preprocessing, stratified cross-validation, dataset splitting, feature standardization, class weight computation, and performance evaluation. Model performance was assessed using several evaluation metrics, including accuracy, precision, recall, F1-score, Matthews correlation coefficient (MCC), area under the receiver operating characteristic curve (AUROC), and area under the precision–recall curve (AUPR).

The proposed framework was evaluated on a workstation running Ubuntu 20.04.6 LTS (64-bit). The hardware configuration consisted of an Intel^®^ Core™ i5-8400 CPU operating at 2.80 GHz, 64 GB DDR4 memory (2133 MHz), a 1 TB hard disk drive (HDD), and an NVIDIA GeForce GTX 1080 graphics processing unit (GPU), which was used to accelerate deep learning model training.

### 5.1 Confusion Matrix

The confusion matrix illustrates the performance of the classification model, displaying the number of correct and incorrect predictions for each class. The diagonal represents accurate classifications, while off-diagonal entries indicate misclassifications between true and predicted labels. Figure 3 shows the confusion matrix of the predicted labels against the true labels of 32 labels ( 31 positive labels plus the Negative interaction label). As the figure shows, most of the labels are predicted correctly.

**Fig 3.**
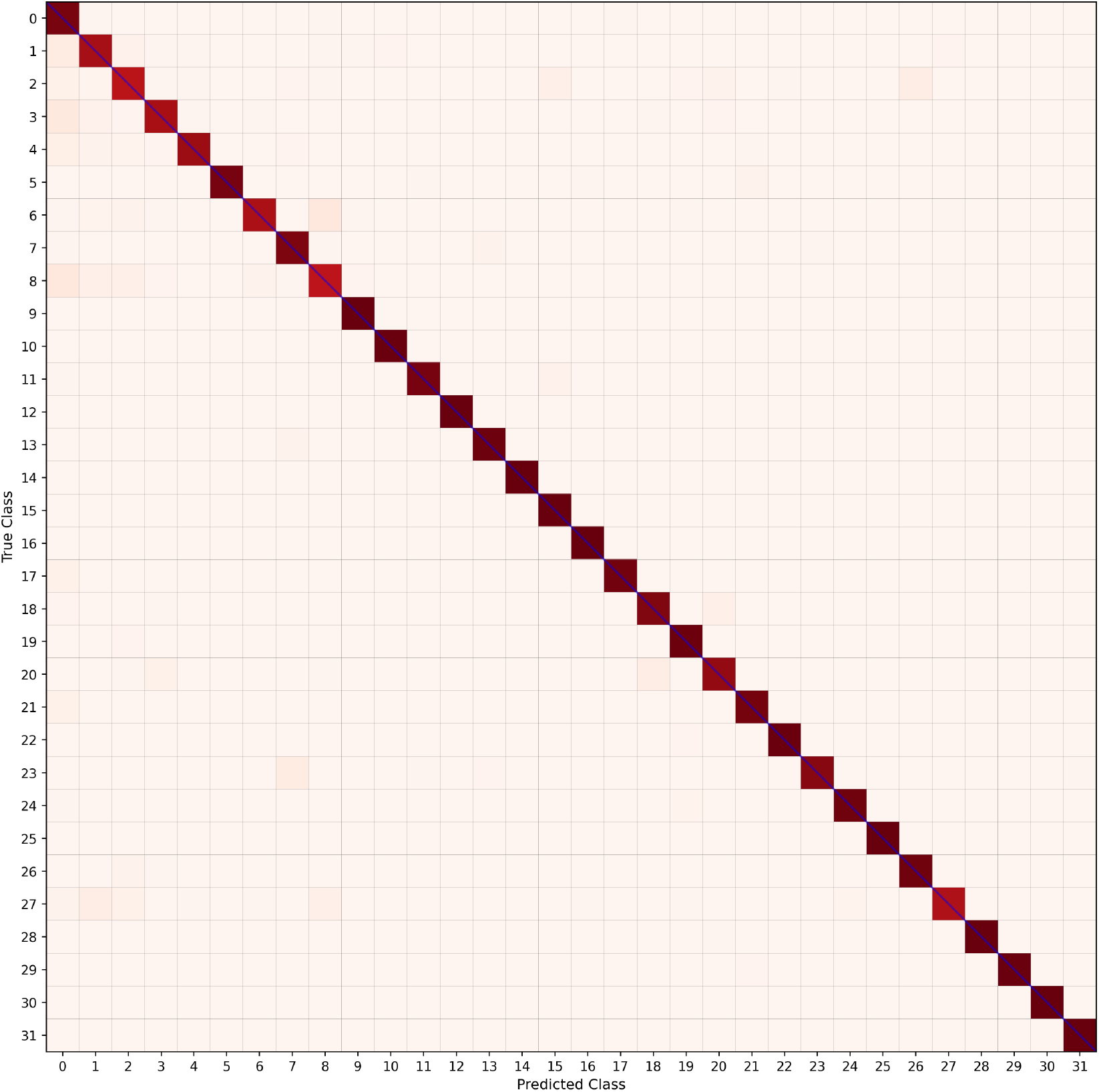
Confusion matrix for the classification model. Diagonal entries show correct predictions, while off-diagonal entries highlight classification errors between true and predicted classes.

### 5.2 Evaluation Protocol

This section evaluates the proposed B-SMART-Former using three complementary evaluation strategies: micro, macro, and weighted-averaged metrics. To provide a comprehensive assessment of the model under different class distribution perspectives, all three evaluation schemes are reported.

#### Micro

Micro-averaged metrics calculate performance by aggregating contributions across all classes. This approach gives equal weight to each individual sample, making it particularly suitable for scenarios where class distribution is imbalanced. Micro-averaging provides a global perspective of model performance by counting total true positives, false negatives, and false positives across all classes [48].

#### Macro

Macro-averaged metrics compute performance for each class independently and then average the results. This method treats all classes equally regardless of their frequency in the dataset. Macro-averaging is especially important for understanding performance on minority classes and ensures that the model performs well across all categories, not just the most common ones [49].

#### Weighted

Weighted-averaged metrics calculate macro-averaged metrics but weight each class’s contribution by its support (the number of true instances for each class). This approach balances the need for class equality with the practical consideration of class distribution. Weighted metrics provide a compromise between micro and macro perspectives, reflecting both overall performance and class distribution0considerations [49].

### 5.3 Comparison with Similarity-Based Methods

Similarity-based methods have consistently achieved outstanding performance in drug–drug interaction prediction by incorporating prior knowledge through similarity matrices [11, 23]. Unlike similarity-independent approaches that rely solely on intrinsic molecular representations, these methods explicitly model relationships between drugs based on chemical structures, protein sequences, biological functions, or known interaction patterns. By integrating this complementary relational information with learned molecular features, similarity-based frameworks are often able to achieve superior predictive performance.

Despite their effectiveness, similarity-based approaches have several practical limitations. Constructing reliable similarity matrices requires additional preprocessing, extensive feature engineering, domain-specific knowledge, and extra computational resources, all of which increase the complexity of the prediction pipeline. Furthermore, when new biotech or small-molecule drugs are introduced, the corresponding similarity matrices must be recomputed and the graph representation updated, which may also require retraining or fine-tuning the predictive model depending on the underlying framework. Consequently, the applicability of similarity-based methods may be limited in dynamic real-world settings where similarity information is incomplete, unavailable, or continuously evolving.

To address these limitations, the proposed B-SMART-Former is designed as a similarity-independent framework that learns directly from intrinsic molecular representations, including ChemBERTa embeddings, ProtBERT embeddings, and molecular fingerprints. By eliminating the dependence on externally computed similarity matrices, the proposed framework substantially simplifies the prediction pipeline while maintaining highly competitive predictive performance.

Table 4 presents a comprehensive comparison between the proposed B-SMART-Former and the state-of-the-art similarity-based framework BSI-Net [11], as a similarity-based approach, under the micro, macro, and weighted-average evaluation regimes. To further investigate the contribution of similarity information, we evaluated a similarity-enhanced version of the proposed framework, denoted B-SMART-Former(+Sim), along with a similarity-free version of BSI-Net, referred to as BSI-Net(-Sim). This comparison enables a direct and fair assessment of both the effectiveness of the proposed architecture and the impact of incorporating similarity information into each framework.

**Table 4.** Performance Comparison of Different Models Across Evaluation Regimes.

| Regime | Metric | B-SMART-Former | B-SMART-Former(+Sim) | BSI-Net(-Sim) [11] | BSI-Net [11] |
| --- | --- | --- | --- | --- | --- |
| Micro | MCC(std) | 0.9013(0.0027) | 0.9556(0.0012) | 0.6239(0.0229) | 0.9809(0.012) |
|  | AUROC(std) | 0.9978(0.0002) | 0.9993(0.0007) | 0.9871(0.0024) | 1.0000(0.000) |
|  | AUPR(std) | 0.9682(0.0024) | 0.9912(0.0019) | 0.4238(0.0182) | 0.9908(0.009) |
| Macro | Precision(std) | 0.8306(0.1725) | 0.9015(0.0127) | 0.4980(0.3775) | 0.9705(0.053) |
|  | Recall(std) | 0.9479(0.0692) | 0.9767(0.0149) | 0.3309(0.3317) | 0.9609(0.073) |
|  | F1-Score(std) | 0.8738(0.1265) | 0.9326(0.0216) | 0.3422(0.3196) | 0.9627(0.058) |
|  | AUROC(std) | 0.9963(0.0081) | 0.9994(0.0071) | 0.9633(0.0283) | 0.9999(0.000) |
|  | AUPR(std) | 0.9311(0.1241) | 0.9843(0.0302) | 0.4238(0.3144) | 0.9908(0.026) |
| Weighted | Precision(std) | 0.9370(0.0734) | 0.9708(0.0025) | 0.7199(0.2051) | 0.9866(0.022) |
|  | Recall(std) | 0.9283(0.0636) | 0.9677(0.0013) | 0.7392(0.3269) | 0.9861(0.025) |
|  | F1-Score(std) | 0.9303(0.0601) | 0.9683(0.0027) | 0.7052(0.2732) | 0.9861(0.021) |
|  | AUROC(std) | 0.9942(0.0042) | 0.9989(0.0041) | 0.9492(0.0266) | 0.9999(0.000) |
|  | AUPR(std) | 0.9724(0.0466) | 0.9932(0.0029) | 0.7550(0.2540) | 0.9981(0.007) |

The results demonstrate that the proposed B-SMART-Former consistently outperforms BSI-Net(-Sim) across all evaluation regimes and performance metrics. Under the micro-averaged evaluation, for example, the proposed model improves the MCC from 0.6239 to 0.9013 and the AUPR from 0.4238 to 0.9682. Similar improvements are consistently observed under the macro and weighted-averaged evaluation regimes, with substantial gains across Precision, Recall, F1-score, AUROC, and AUPR. Notably, the macro-averaged F1-score increases from 0.3422 to 0.8738, while the weighted-averaged F1-score improves from 0.7052 to 0.9303. These results demonstrate that the proposed Transformer–ResNet architecture effectively learns highly discriminative molecular representations directly from ChemBERTa embeddings, ProtBERT embeddings, and molecular fingerprints without relying on externally computed similarity information.

The inclusion of similarity matrices further enhances the predictive capability of the proposed framework. Compared with the similarity-free version, B-SMART-Former(+Sim) improves the micro-averaged MCC from 0.9013 to 0.9556, the AUROC from 0.9978 to 0.9993, and the AUPR from 0.9682 to 0.9912. Similar improvements are consistently observed under the macro and weighted-averaged evaluation regimes, indicating that similarity information provides complementary relational knowledge beyond intrinsic molecular representations. Nevertheless, the relatively small performance gap between B-SMART-Former and B-SMART-Former(+Sim) indicates that the proposed architecture is already capable of learning highly informative molecular representations directly from intrinsic drug features, while similarity information provides an additional but incremental performance improvement.

Although BSI-Net achieves the highest performance on most evaluation metrics, B-SMART-Former(+Sim) remains highly competitive across all evaluation regimes and even slightly surpasses BSI-Net in terms of the micro-averaged AUPR (0.9912 vs. 0.9908). Similar trends are observed under the macro and weighted-averaged evaluation regimes, where the differences between the two methods remain consistently small. Moreover, the proposed framework exhibits low standard deviations across the 10-fold stratified cross-validation, indicating stable and robust predictive performance.

A key advantage of the proposed framework is that it eliminates the dependence on externally computed similarity matrices while maintaining highly competitive predictive performance. Unlike graph-based approaches that require additional preprocessing, feature engineering, and similarity computation, B-SMART-Former learns directly from ChemBERTa embeddings, ProtBERT embeddings, and molecular fingerprints. This substantially simplifies the prediction pipeline and makes the framework more readily applicable to newly introduced biotech and small-molecule drugs for which reliable similarity information may not yet be available.

Overall, these results demonstrate that B-SMART-Former effectively narrows the performance gap between similarity-free and similarity-based approaches. Even without handcrafted similarity information, the proposed architecture substantially outperforms BSI-Net(-Sim), while its similarity-enhanced version approaches—and in selected metrics slightly exceeds—the performance of the full BSI-Net. These findings highlight the effectiveness, robustness, and practical applicability of the proposed framework for multiclass biotech–small-molecule drug–drug interaction prediction.

### 5.4 Micro-averaged Performance

Table 5 reports the micro-averaged performance of all compared methods. Among the evaluated models, B-SMART-Former achieves the highest MCC, AUROC, and AUPR values, demonstrating superior overall predictive performance under severe class imbalance.

**Table 5.** Performance comparison of various methods in the micro regime. Bold values indicate the best performance for each metric. Standard deviations are shown in parentheses.

| Methods | MCC(std) | AUROC(std) | AUPR(std) |
| --- | --- | --- | --- |
| SVM [51] | 0.8927(0.0022) | 0.9968(0.0002) | 0.9624(0.0019) |
| XGBoost [52] | 0.8305(0.0037) | 0.9981(0.0001) | 0.9545(0.0020) |
| MLP [11] | 0.6564(0.0062) | 0.9881(0.0006) | 0.7577(0.0130) |
| CNN [50] | 0.1794(0.0623) | 0.8541(0.0629) | 0.2181(0.0638) |
| B-SMART-Former | **0.9013(0.0027)** | **0.9978(0.0002)** | **0.9682(0.0024)** |

Compared with conventional machine learning methods and deep learning baselines, particularly CNN [50], the proposed architecture consistently achieves better discrimination between interaction classes. The strong performance can be attributed to the complementary strengths of Transformer-based global dependency modeling and residual convolutional feature extraction, which together produce highly informative representations for multiclass drug–drug interaction prediction.

### 5.5 Macro-averaged Performance

Table 6 presents the macro-averaged performance of all compared methods. The proposed B-SMART-Former achieves the highest Recall, F1-score, AUROC, and AUPR among all baseline methods, while XGBoost achieves the highest Precision.

**Table 6.** Performance comparison of various methods in the macro regime. Bold values indicate the best performance for each metric. Standard deviations are shown in parentheses.

| Methods | Precision(std) | Recall(std) | F1-Score(std) | AUROC(std) | AUPR(std) |
| --- | --- | --- | --- | --- | --- |
| SVM [51] | 0.8659(0.0065) | 0.8653(0.0050) | 0.8610(0.0035) | 0.9923(0.0010) | 0.9059(0.0046) |
| XGBoost [52] | **0.9286(0.0049)** | 0.7792(0.0116) | 0.8296(0.0096) | 0.9958(0.0015) | 0.9237(0.0057) |
| MLP [11] | 0.5537(0.0100) | 0.9267(0.0065) | 0.6472(0.0091) | 0.9921(0.0010) | 0.8572(0.0106) |
| CNN [50] | 0.3570(0.0427) | 0.5113(0.1258) | 0.3152(0.0637) | 0.8835(0.0610) | 0.4160(0.1071) |
| B-SMART-Former | 0.8306(0.1725) | **0.9479(0.0692)** | **0.8738(0.1265)** | **0.9963(0.0081)** | **0.9311(0.1241)** |

The superior macro-averaged performance demonstrates that the proposed architecture generalizes well across both frequent and infrequent interaction classes without excessively favoring the dominant classes. In particular, the improvements in Recall and F1-score indicate that B-SMART-Former effectively captures minority interaction patterns while maintaining stable overall classification performance.

### 5.6 Weighted-averaged Performance

Table 7 summarizes the weighted-averaged performance of all compared methods. B-SMART-Former consistently achieves the highest scores across all evaluation metrics, confirming its robustness under realistic class distributions where different interaction types occur with varying frequencies.

**Table 7.** Performance comparison of various methods in the weighted regime. Bold values indicate the best performance for each metric. Standard deviations are shown in parentheses.

| Methods | Precision(std) | Recall(std) | F1-Score(std) | AUROC(std) | AUPR(std) |
| --- | --- | --- | --- | --- | --- |
| SVM [51] | 0.9256(0.0016) | 0.9210(0.0017) | 0.9222(0.0016) | 0.9926(0.0007) | 0.9609(0.0021) |
| XGBoost [52] | 0.8829(0.0025) | 0.8797(0.0026) | 0.8713(0.0029) | 0.9878(0.0006) | 0.9495(0.0021) |
| MLP [11] | 0.8366(0.0037) | 0.6936(0.0081) | 0.7175(0.0091) | 0.9706(0.0022) | 0.9056(0.0024) |
| CNN [50] | 0.6468(0.0597) | 0.1694(0.0674) | 0.1736(0.0726) | 0.7381(0.0743) | 0.4942(0.0756) |
| B-SMART-Former | **0.9370(0.0734)** | **0.9283(0.0636)** | **0.9303(0.0601)** | **0.9942(0.0042)** | **0.9724(0.0466)** |

Compared with the competing approaches, the proposed model provides a better balance between precision and recall while maintaining excellent discriminative capability, as reflected by its AUROC and AUPR values. These findings demonstrate that the proposed architecture is well suited for practical multiclass drug–drug interaction prediction tasks involving highly imbalanced datasets.

Figure 4 presents the AUROC and AUPR values achieved by the proposed B-SMART-Former across the 32 interaction classes. Mean values and standard deviations are reported for the micro, macro, and weighted averaging schemes.

**Fig 4.**
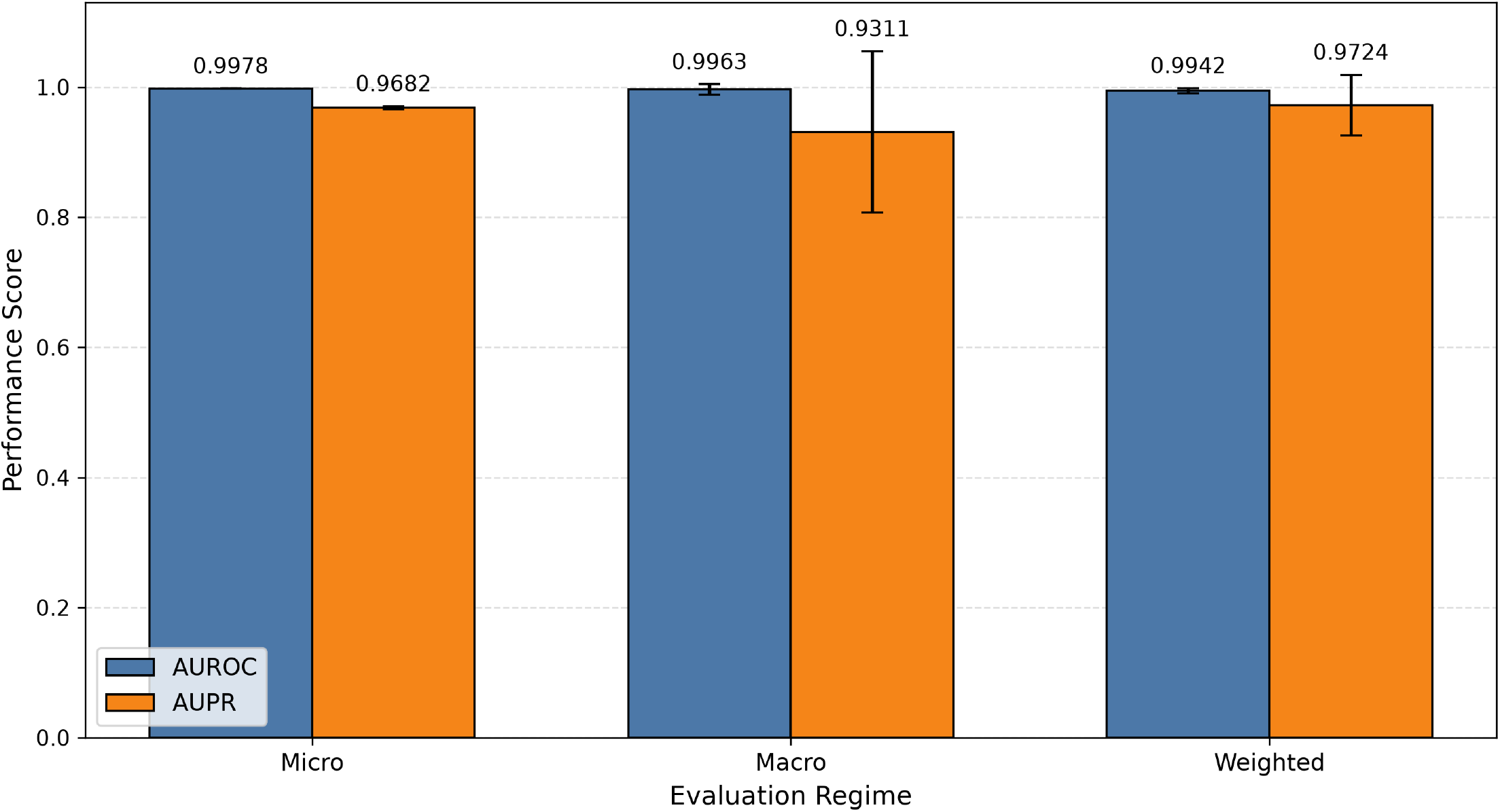
AUROC and AUPR results of B-SMART-Former. The mean and standard deviation across 10-fold stratified cross-validation are reported for the micro, macro, and weighted averaging schemes.

### 5.7 Ablation Study

To systematically assess the contribution of each component in the proposed B-SMART-Former architecture, we conducted an ablation study by progressively removing or modifying individual modules while keeping all other architectural configurations, hyperparameters, and training settings unchanged. As described in the previous sections, B-SMART-Former integrates three principal components: a Transformer (TransF) module [33], a Residual Network (ResNet) module [41], and a Multi-Layer Perceptron (MLP) module [11]. The effectiveness of each component was evaluated both independently and in combination with the remaining modules, and the resulting model variants were compared against the complete B-SMART-Former architecture to quantify the contribution of each module to the overall predictive performance.

#### Micro

Table 8 summarizes the performance of the resulting model variants using micro-averaged evaluation metrics. The performance of different model variants is compared using micro-averaged MCC, AUROC, and AUPR, to assess the contribution of each architectural component to the overall predictive performance.

**Table 8.**
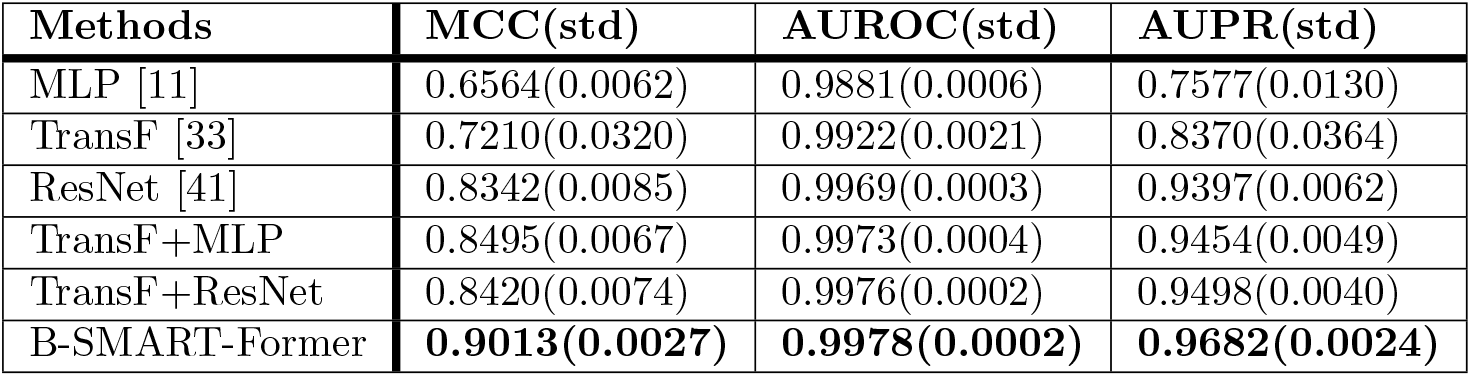
Ablation study of B-SMART-Former based on micro-averaged evaluation metrics.

#### Macro

Table 9 presents the corresponding macro-averaged evaluation results, providing a balanced assessment of the model performance across all interaction classes by assigning equal importance to each class. The performance of different model variants is evaluated using macro-averaged Precision, Recall, F1-score, AUROC, and AUPR, highlighting the impact of each model component while assigning equal importance to all interaction classes.

**Table 9.**
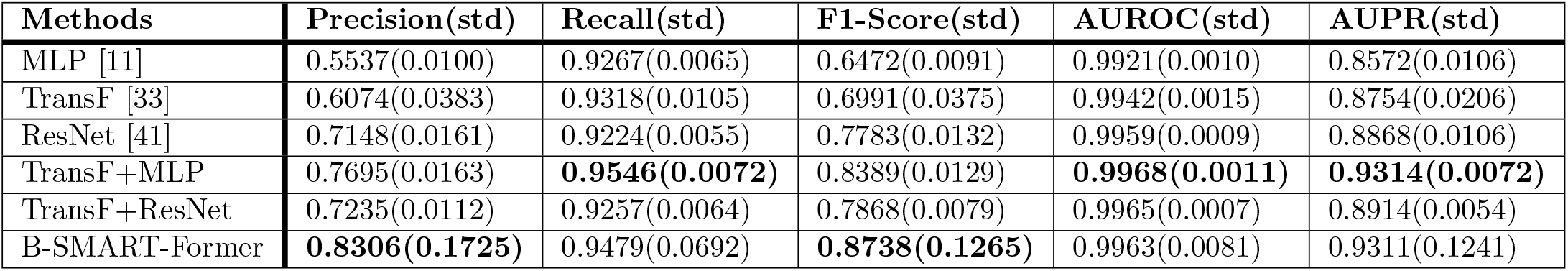
Ablation study of B-SMART-Former based on macro-averaged evaluation metrics.

| Methods | Precision(std) | Recall(std) | F1-Score(std) | AUROC(std) | AUPR(std) |
| --- | --- | --- | --- | --- | --- |
| MLP [11] | 0.5537(0.0100) | 0.9267(0.0065) | 0.6472(0.0091) | 0.9921(0.0010) | 0.8572(0.0106) |
| TransF [33] | 0.6074(0.0383) | 0.9318(0.0105) | 0.6991(0.0375) | 0.9942(0.0015) | 0.8754(0.0206) |
| ResNet [41] | 0.7148(0.0161) | 0.9224(0.0055) | 0.7783(0.0132) | 0.9959(0.0009) | 0.8868(0.0106) |
| TransF+MLP | 0.7695(0.0163) | **0.9546(0.0072)** | 0.8389(0.0129) | **0.9968(0.0011)** | **0.9314(0.0072)** |
| TransF+ResNet | 0.7235(0.0112) | 0.9257(0.0064) | 0.7868(0.0079) | 0.9965(0.0007) | 0.8914(0.0054) |
| B-SMART-Former | **0.8306(0.1725)** | 0.9479(0.0692) | **0.8738(0.1265)** | 0.9963(0.0081) | 0.9311(0.1241) |

#### Weighted

Table 10 reports the weighted-averaged evaluation results, where the contribution of each interaction class is proportional to its frequency in the dataset, offering an overall performance assessment under the original class distribution. The performance of different model variants is compared using weighted-averaged Precision, Recall, F1-score, AUROC, and AUPR, where each interaction class contributes proportionally according to its frequency in the dataset.

**Table 10.**
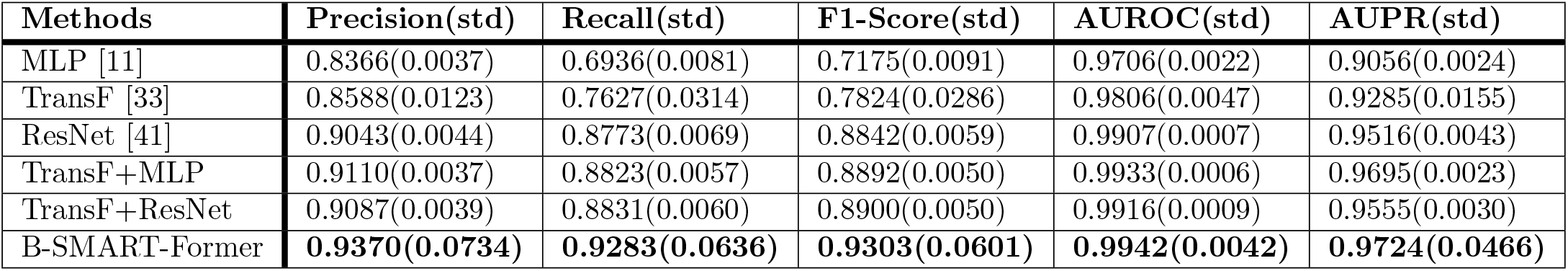
Ablation study of B-SMART-Former based on weighted-averaged evaluation metrics.

### 5.8 XAI Results and Interpretation

Figure 5 presents the global attribution analysis obtained using Integrated Gradients. Attribution scores were aggregated across all explained samples and grouped according to the three feature modalities used in the proposed framework: ChemBERTa embeddings, molecular fingerprint features, and ProtBERT-derived biotech embeddings.

**Fig 5.**
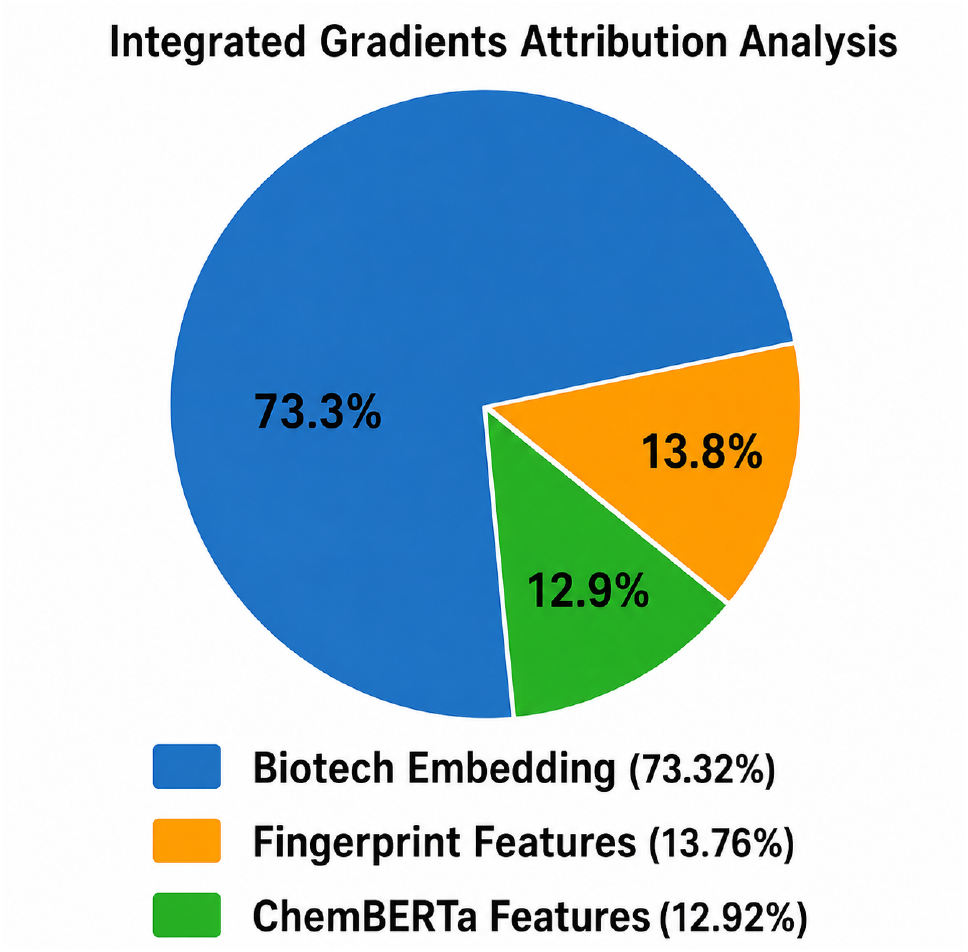
Global feature-group attribution obtained using Integrated Gradients. Attribution scores were aggregated across all explained samples and normalized to percentage contributions. Biotech (ProtBERT) features contributed 73.3% of the total attribution score, while molecular fingerprints and ChemBERTa features contributed 13.8% and 12.9%, respectively.

The results show that biotech features contributed the largest proportion of the overall attribution score, accounting for approximately 73.3% of the total contribution. In comparison, molecular fingerprint features and ChemBERTa embeddings contributed 13.8% and 12.9%, respectively. These findings indicate that the proposed framework relies primarily on information derived from biotech drugs while also incorporating complementary information from small-molecule representations.

The similar contributions of ChemBERTa embeddings and molecular fingerprints suggest that both learned molecular representations and structural descriptors provide useful information for interaction prediction. At the same time, the dominant contribution of ProtBERT features highlights the importance of biotech-drug sequence information in determining interaction outcomes. Overall, the attribution analysis demonstrates that the model integrates information from multiple feature sources rather than relying exclusively on a single modality.

Representative local explanations generated using Integrated Gradients are shown in Figure 6. The highlighted atoms correspond to molecular regions that received the highest attribution scores and therefore had the greatest influence on the model’s predictions. Across different compounds, the model consistently focused on specific structural motifs, including heterocyclic rings, aromatic groups, carbonyl-containing regions, amide groups, and oxygen-containing linkers.

**Fig 6.**
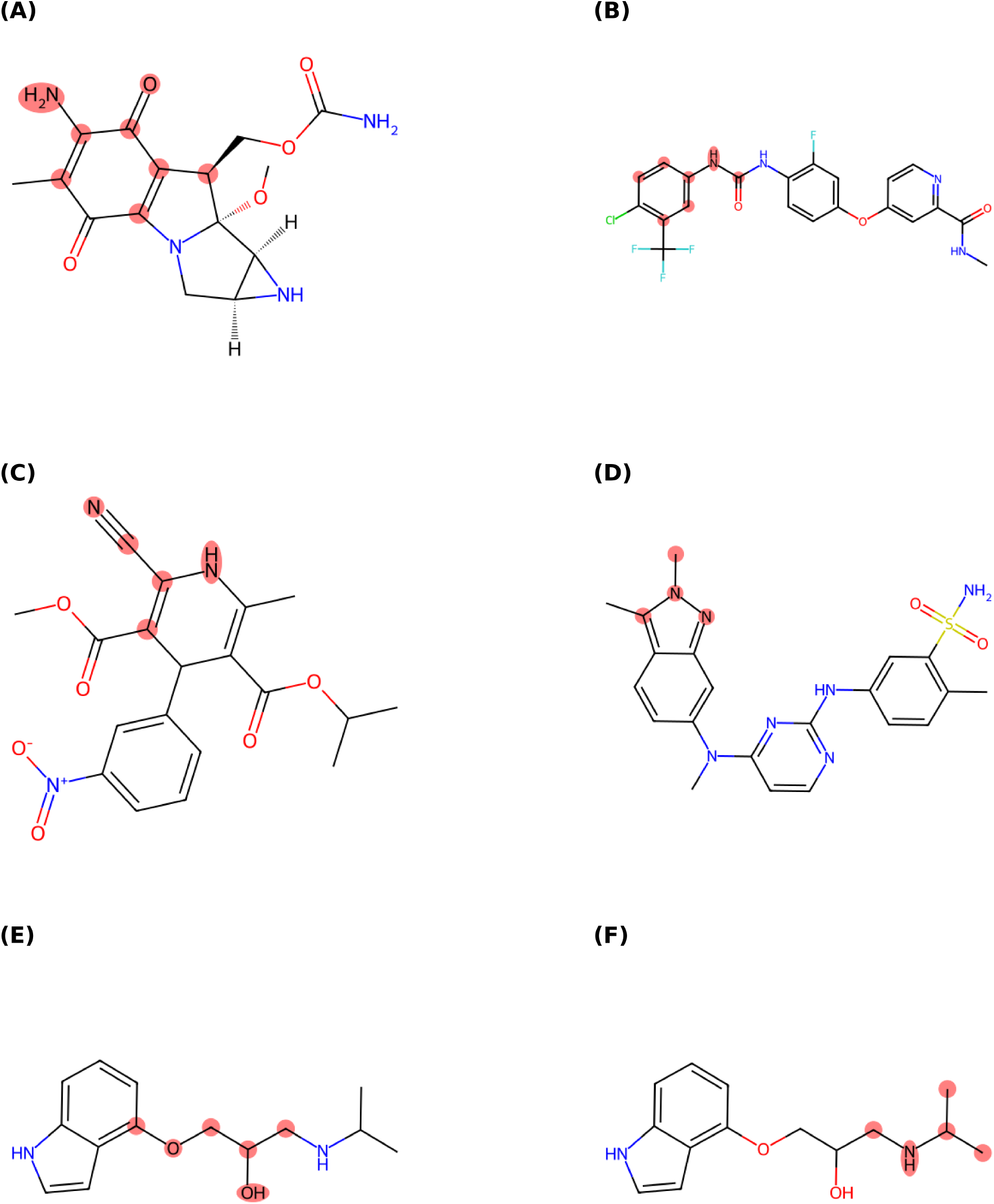
Representative molecular explanations generated using Integrated Gradients. Highlighted atoms indicate molecular regions with the highest attribution scores and therefore the strongest influence on model predictions.

These results provide additional insight into the decision-making process of the proposed framework by identifying the molecular regions that contributed most strongly to individual predictions. Although Integrated Gradients does not establish causal biochemical mechanisms, the consistency of the highlighted motifs across multiple compounds suggests that the model captures meaningful structural patterns associated with biotech–small-molecule interactions.

Table 11 summarizes representative molecular substructures identified within the highlighted regions shown in Figure 6. These examples illustrate the types of structural motifs that were repeatedly assigned high attribution scores by the model. The recurrence of such motifs across different compounds further supports the interpretability of the proposed framework and provides a qualitative view of the molecular patterns utilized during prediction.

**Table 11.** Representative molecular substructures highlighted by Integrated Gradients.

| Figure | Highlighted region |
| --- | --- |
| (A) | Carbonyl-containing heterocyclic scaffold |
| (B) | Halogenated aromatic ring linked to a urea group |
| (C) | Cyano-substituted heterocycle |
| (D) | Nitrogen-rich fused aromatic system |
| (E) | Ether linkage and hydroxyl-containing side chain |
| (F) | Amide-containing terminal region |

## 6 Discussion and Conclusion

This work introduced B-SMART-Former, a hybrid Transformer-based framework for predicting drug–drug interactions between biotech and small-molecule drugs. To improve both generalization and feature representation, the proposed framework leverages foundation models by combining ChemBERTa embeddings and molecular fingerprints for small molecules with ProtBERT embeddings for biotech drugs. These complementary representations are integrated and processed through a Transformer architecture followed by residual convolutional networks to learn informative joint embeddings for interaction prediction. Experimental results demonstrate that B-SMART-Former achieves performance comparable to similarity-based approaches despite relying solely on intrinsic molecular representations. Unlike similarity-based methods, the proposed framework does not require prior knowledge of relationships among drugs, making it more suitable for predicting interactions involving previously unseen drugs.

To improve the interpretability of the proposed framework, we applied the post-hoc explainability method Integrated Gradients to identify the molecular features that contribute most to the predicted interactions. The resulting attribution maps demonstrate the influence of specific chemical substructures on the model’s predictions and provide valuable insights into its decision-making process. To the best of our knowledge, this is the first study to investigate post-hoc explainability for biotech–small-molecule drug–drug interaction prediction.

Future work will focus on developing more expressive architectures that capture complex dependencies between molecular representations while preserving the ability to generalize without relying on similarity-based features. Another important direction is to improve the robustness of prediction models against noisy or incomplete input representations. Finally, incorporating more advanced explainability techniques and validating the generated explanations with domain experts will further enhance the transparency, reliability, and practical applicability of deep learning models for biotech–small-molecule drug interaction prediction.

## 7 Data Availability

The source code and implementation of B-SMART-Former are publicly available at: https://github.com/BioinformaticsIASBS/B-SMART-Former.

## 8 Supporting information

### Abbreviations

This appendix summarizes the mathematical notation used throughout the proposed framework. For clarity, the notation is organized according to the main stages of the methodology. Table 12 lists the symbols used in the feature representation stage. Table 13 summarizes the notation associated with the proposed B-SMART-Former architecture, including the input projection layers, Transformer module, residual convolutional module, and classification head. Finally, Table 14 presents the notation used in the Integrated Gradients-based explainability analysis.

**Table 12.** Notation used in the feature representation module.

| Symbol | Description |
| --- | --- |
| $i$ | Index of the small-molecule drug |
| $j$ | Index of the biotech drug |
| $t$ | Index of a SMILES token |
| $k$ | Index of an amino acid residue |
| $\ell$ | Length of the tokenized SMILES sequence |
| $\ell_P$ | Length of the protein sequence |
| $S$ | Tokenized SMILES sequence of a small-molecule drug |
| $P$ | Amino acid sequence of a biotech drug |
| $s_t$ | $t$-th SMILES token |
| $p_k$ | $k$-th amino acid residue |
| $E(\cdot)$ | Token embedding function |
| $P_t$ | Positional embedding of the $t$-th SMILES token |
| $\mathbf{x}_t$ | Input embedding of the $t$-th SMILES token |
| $\mathbf{H}$ | Contextual hidden-state matrix generated by ChemBERTa |
| $\mathbf{h}_t$ | Contextual representation of the $t$-th SMILES token |
| $\mathbf{e}^{\text{ChemBERTa}}$ | Mean-pooled ChemBERTa embedding |
| $G = (V, E)$ | Molecular graph of a small-molecule drug |
| $V$ | Set of atoms in the molecular graph |
| $E$ | Set of chemical bonds in the molecular graph |
| $\Phi(\cdot)$ | Morgan fingerprint generation procedure |
| $r$ | Radius parameter of the Morgan fingerprint |
| $d_{\text{FP}}$ | Morgan fingerprint dimensionality (number of bits) |
| $\mathbf{f}_{\text{FP}}$ | Binary Morgan fingerprint vector |
| $\mathbf{x}_{SM}^{(i)}$ | Final representation of the $i$-th small-molecule drug |
| $\mathbf{Z}$ | Contextual hidden-state matrix generated by ProtBERT |
| $\mathbf{z}_k$ | Contextual representation of the $k$-th amino acid residue |
| $\mathbf{x}_{BT}^{(j)}$ | Final representation of the $j$-th biotech drug |
| $\oplus$ | Vector concatenation operator |
| $\mathbf{f}_{ij}$ | Final multimodal feature vector of the $(i, j)$ drug pair |
| $\mathbb{R}$ | Set of real numbers |

**Table 13.** Notation used in the proposed B-SMART-Former architecture.

| Symbol | Description |
| --- | --- |
| $\mathbf{h}^{(1)}$ | Output of the first input projection layer |
| $\mathbf{h}^{(2)}$ | Output of the second input projection layer |
| $\mathbf{W}_1$ | Weight matrix of the first projection layer |
| $\mathbf{W}_2$ | Weight matrix of the second projection layer |
| $\mathbf{b}_1$ | Bias vector of the first projection layer |
| $\mathbf{b}_2$ | Bias vector of the second projection layer |
| $\text{BN}(\cdot)$ | Batch normalization operation |
| $\text{Dropout}(\cdot)$ | Dropout operation |
| $\text{Swish}(\cdot)$ | Swish activation function |
| $\mathbf{X}$ | Input token sequence to the Transformer encoder |
| $L$ | Length of the Transformer input sequence |
| $d$ | Embedding dimension of each Transformer token |
| $\mathbf{Q}$ | Query matrix |
| $\mathbf{K}$ | Key matrix |
| $\mathbf{V}$ | Value matrix |
| $\mathbf{W}_Q$ | Query projection matrix |
| $\mathbf{W}_K$ | Key projection matrix |
| $\mathbf{W}_V$ | Value projection matrix |
| $d_k$ | Dimensionality of the key vectors |
| $\text{Attention}(\cdot)$ | Scaled dot-product self-attention |
| $\text{MHA}(\cdot)$ | Multi-head self-attention |
| $\text{head}_i$ | Output of the $i$-th attention head |
| $H$ | Number of attention heads |
| $\mathbf{W}_O$ | Output projection matrix |
| $\mathbf{T}$ | Transformer output feature sequence |
| $\mathbf{C}$ | Output feature map of the convolutional layer |
| $\mathbf{R}$ | Output of the residual convolutional block |
| $\text{Conv1D}(\cdot)$ | One-dimensional convolution |
| $\mathcal{F}(\cdot)$ | Residual nonlinear transformation |
| $\mathbf{g}$ | Fused representation before classification |
| $\mathbf{g}^{\text{Trans}}$ | Transformer branch representation |
| $\mathbf{g}^{\text{Res}}$ | Residual branch representation |
| $\mathbf{o}^{(l)}$ | Hidden representation of the $l$-th MLP layer |
| $\mathbf{W}^{(l)}$ | Weight matrix of the $l$-th MLP layer |
| $\mathbf{b}^{(l)}$ | Bias vector of the $l$-th MLP layer |
| $\hat{\mathbf{y}}$ | Predicted probability distribution |
| $\mathbf{W}_o$ | Output-layer weight matrix |
| $\mathbf{b}_o$ | Output-layer bias vector |
| $\text{Softmax}(\cdot)$ | Softmax activation function |

**Table 14.** Notation used in the explainability analysis.

| Symbol | Description |
| --- | --- |
| $IG_i(x)$ | Integrated Gradients attribution of the $i$-th feature |
| $x$ | Input sample to be explained |
| $x'$ | Baseline input (zero vector) |
| $\alpha$ | Interpolation coefficient |
| $F(\cdot)$ | Prediction function of the trained model |
| $M$ | Trained B-SMART-Former model |
| $X_{\text{test}}$ | Set of selected test samples for explanation |
| $\mathbf{A}_x$ | Integrated Gradients attribution vector |
| IntegratedGradients(·) | Integrated Gradients method |

